# Identification of the cell-type-specific ER membrane protein Tanmp expressed in hypothalamic tanycytes and subsets of neurons

**DOI:** 10.1101/2021.07.06.451253

**Authors:** Osamu Takahashi, Mayuko Tanahashi, Saori Yokoi, Mari Kaneko, Tomoko Tokuhara, Kaori Yanaka, Shinichi Nakagawa, Hiroshi Maita

## Abstract

Genomes of higher eukaryotes encode many uncharacterized proteins, and the functions of these proteins cannot be predicted from the primary sequences due to a lack of conserved functional domains. During a screening of novel noncoding RNAs abundantly expressed in mouse brains, we incidentally identified a gene termed Tanmp, which encoded an endoplasmic reticulum (ER) protein without known functional domains. Tanmp is specifically expressed in the nervous system, and the highest expression was observed in a specialized cell type called tanycyte that aligns the ventral wall of the third ventricle in the hypothalamus. Immunostaining of Tanmp revealed the fine morphology of tanycytes with highly branched apical ER membranes. Immunoprecipitation revealed that Tanmp associates with mitochondrial ATPase at least *in vitro*, and ER and mitochondrial signals occasionally overlapped in tanycytes. Mutant mice lacking Tanmp did not exhibit overt phenotypes, suggesting that Tanmp is not essential in mice reared under normal laboratory conditions. We also found that RNA probes that are predicted to uniquely detect Tanmp mRNA cross-reacted with uncharacterized RNAs, highlighting the importance of experimental validation of the specificity of probes during the hybridization-based study of RNA localization.

## Introduction

Transcripts longer than 200 nucleotides that lack protein coding capacity are called long noncoding RNAs (lncRNAs), of which the threshold was arbitrarily set during early studies done in the 2000s (reviewed in Ulitsky, 2016). Meanwhile, the development of ribosome profiling technology has revealed that many of the transcripts that were initially described as lncRNAs indeed associate with ribosomes (Ingolia, 2014). These transcripts are believed to be translated into peptides, which are shorter than the arbitrarily set thresholds for the lack of protein coding capacity of lncRNAs. A series of studies using mutant animals revealed that novel proteins or peptides encoded by “lncRNAs” play important physiological roles, including the regulation of transcription via inhibition of P-TEF-b (Hanyu-Nakamura et al., 2008), activation of G protein-coupled receptors (Chng et al., 2013; Pauli et al., 2014), proteolytic processing of transcription factors (Kondo et al., 2007), modulation of Ca^2+^ pump activity (Anderson et al., 2015; Nelson et al., 2016), regulation of mTORC1 (Matsumoto et al., 2017), and regulation of mRNA de-capping complexes (D’Lima et al., 2017). These newly discovered peptides lack known conserved domains and thus have long been overlooked by functional predictions based on primary sequence analyses. Studies have also shown that a group of novel proteins, entirely consisting of intrinsically disordered regions, possesses outstanding molecular properties to inhibit the formation of disease-related molecular aggregates or protect DNA from irradiation (Hashimoto et al., 2016; Tsuboyama et al., 2020). Interestingly, at least two of these “superdisordered” proteins maintain their molecular activity even after the random shuffling of their primary amino acid sequences (Tsuboyama et al., 2020), suggesting that the function of these superdisordered proteins is dependent on the composition of the amino acids rather than their primary sequences. The functional independence of the amino acid sequences makes it difficult to predict the function of these proteins by conventional bioinformatic analyses. As such, even 20 years after the first draft sequences of the human and mouse genomes were published, nearly a thousand proteins remain uncharacterized (Frankish et al., 2021). Together with the substantial number of uncharacterized lncRNAs transcribed from the genome, these unpredictable proteins constitute the “dark matter” of the genome, which may regulate important processes in higher organisms.

In this study, we examined the expression and subcellular distribution of the poorly characterized gene *6330403K07Rik/UGS148*, which was previously reported as a specific molecular marker for tanycytes, in detail (Ma et al., 2015). Tanycytes are specialized cell types that align the ventral region of the third ventricle around the medial eminence and are believed to be involved in the transport of small molecules and peptides from the cerebrospinal fluid to neurons in the arcuate nucleus and vice versa (reviewed in Langlet, 2014; Rodriguez et al., 2019). We named this gene Tanmp (tanycyte-associated ER <u>m</u>embrane protein) due to the intense expression in tanycytes and specific subcellular localization in the endoplasmic reticulum (ER) membrane. Immunoprecipitation analyses revealed that Tanmp associates with mitochondrial ATP synthetase at least *in vitro*, suggesting that Tanmp might be involved in the interaction of the ER and the mitochondrial membrane in this specialized cell type. Tanmp knockout (KO) mice were viable and exhibited no overt phenotype, suggesting that Tanmp is not required for the mice, at least under normal laboratory conditions.

## Results

### Incidental identification of Tanmp while identifying novel transcripts that exhibit unique subcellular distribution

To identify novel functional lncRNAs, we initially investigated the subcellular distribution of transcripts of uncharacterized genes by *in situ* hybridization, especially focusing on genes with the suffix -Rik identified by the FANTOM project (Carninci et al., 2005). We expected that if the transcripts are highly enriched in the nucleus, they might be good candidates for a novel functional lncRNA considering that no translation occurs in the nucleus. We used cultured hippocampal neurons for this study because the nervous system generally expresses a broader range of lncRNAs than other tissue and cell types (reviewed in Ng et al., 2013). We arbitrarily selected 24 genes with the suffix “Rik” from the list of highly expressed genes identified by microarray analyses of cultured hippocampal neurons (Supplemental Table T1), and intense signals were confirmed for 16 of them by fluorescent *in situ* hybridization using cultured hippocampal neurons (Fig. 1A). Among them, transcripts of 2 genes, *2610002J23Rik* and *6330403K07Rik*, were highly enriched in the nucleus (Fig. 1A). We focused on *6330403K07Rik* in subsequent studies because of its intense and specific expression in the nervous system.

**Figure 1.**
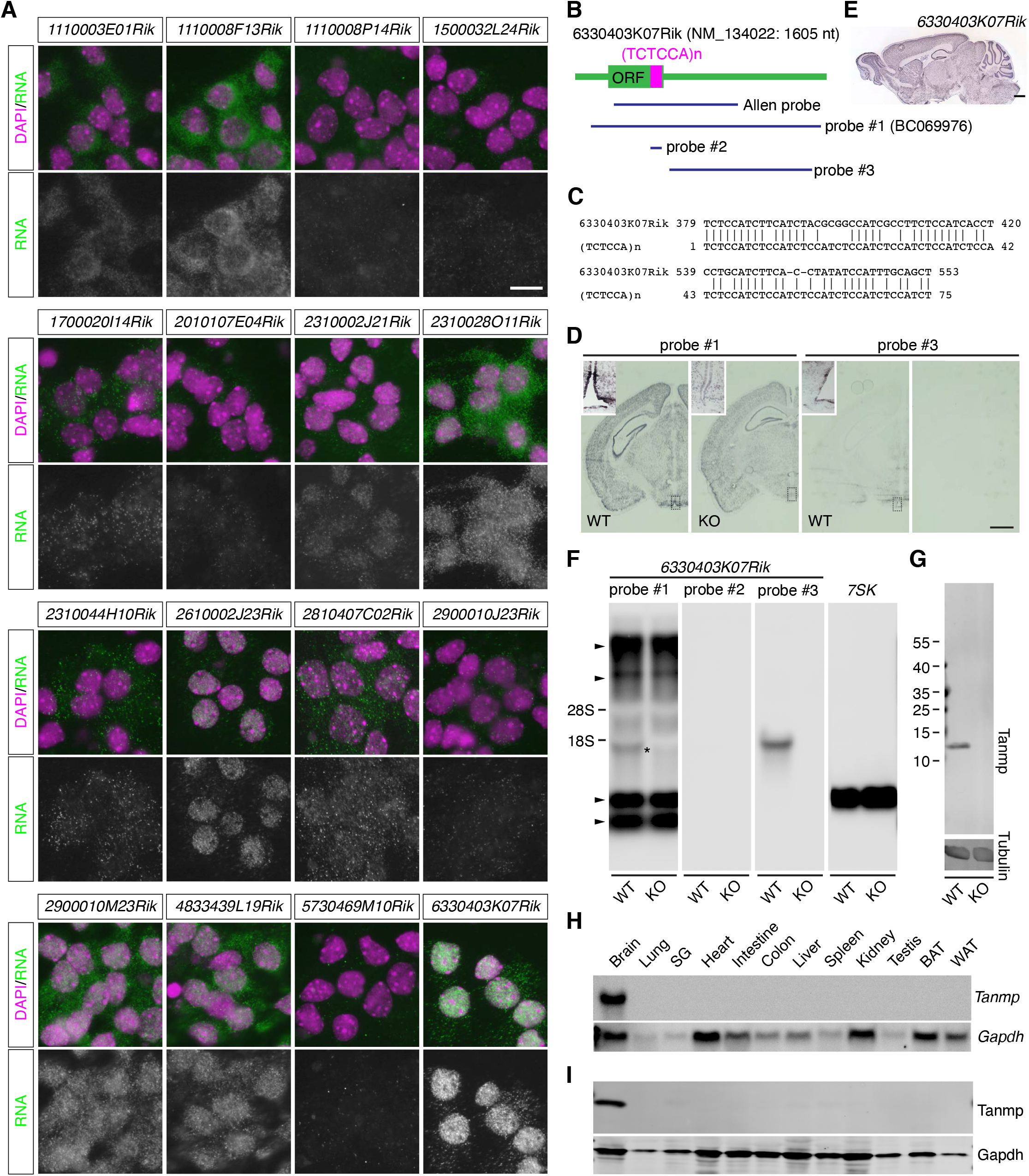
Cross-reactivity of RNA probes and identification of Tanmp using a specific probe. (A) Localization of abundantly expressed transcripts of uncharacterized genes with the -Rik suffix (green). Nuclei were simultaneously detected with DAPI (magenta). (B) Schematic drawing of 6330403K07Rik mRNA and positions of RNA probes. (C) Sequence alignment of the region that exhibits weak homology to (TCTCCA)n revealed by the RepeatMasker search. (D) Coronal sections at the hypothalamus levels showing the expression of 6330403K07Rik in the adult brain detected with the RNA probes shown in (B). Note that probe #1 detected signals in the knockout (KO) mice lacking the expression of this gene. Intense signals in the ventral region of the hypothalamus (inset) were observed only in the brains of the wild-type (WT) mice but not in the brains of the KO mice. (E) Sagittal section of the brain in the Allen Brain Atlas database showing the expression of 6330403K07Rik. The ubiquitous expression was similar to the pattern obtained with probe #1 shown in (B). (F) Northern blot analyses of transcripts detected with probes #1, #2, and #3 shown in (B). Detection of 7SK was performed to confirm equal loading of the RNA samples in each lane. The asterisk shows the band corresponding to the size of 6330403K07Rik mRNA. Arrowheads indicate bands detected in the samples obtained from the WT mice but not in the KO mice. (G) Western blot analyses of Tanmp in the adult mouse brain. Specific bands were detected in the samples prepared from the WT adult brains. (H, I) Tissue distribution of Tanmp mRNA (H) and protein (I) detected by Northern blot and Western blot, respectively. Note that both Tanmp mRNA and protein are specifically expressed in the brain. SG: salivary gland, BAT: brown adipose tissue, WAT: white adipose tissue. Scale bars, 10 μm in A and 1 mm in E and D.

To investigate the expression pattern of *6330403K07Rik* in the adult brain by *in situ* hybridization, we initially used a full-length cDNA clone, BC069976, for preparation of the probe (Tanmp #1, Fig. 1B). The clone did not contain any homologous sequence in the genome when searched by NCBI BLAST and RepeatMasker, except for a short 73 bp fragment that exhibited only weak homology to a simple repeat, (TCTCCA)n (Fig. 1B, C). *In situ* hybridization using a stringent hybridization condition revealed that *6330403K07Rik* was broadly expressed throughout the entire brain regions (probe #1, Fig. 1D), which was consistent with the data in the Allen mouse brain atlas (Fig. 1E), a public genome-wide database for the expression of genes in the mouse brain (Sunkin et al., 2013). However, during this study, a paper reported that *6330403K07Rik/UGS148* is highly enriched in tanycytes, a group of ependymal cells that align ventral regions of the 3^rd^ ventricle in the hypothalamus (Ma et al., 2015). We thus speculated that the probe used in our initial study and the Allen Brain Atlas cross-reacted with additional transcripts via the region weakly homologous to (TCTCCA)n. To test this hypothesis, we prepared probes #2 and #3 that do and do not contain the repeat-homologous region (Fig. 1B) and performed Northern blot analyses using brain RNAs (Fig. 1F). As expected, the probe #1 we initially used detected multiple transcripts (arrowheads in Fig. 1F) in addition to the transcript at the expected size (asterisk in Fig. 1F). To further confirm the cross-reactivity of the probes, we performed Northern blot analyses using brain RNAs derived from mutant mice lacking the *6330403K07Rik/UGS148* transcripts, which were generated during this study. Extra bands were also detected in the mice lacking the *6330403K07Rik/UGS148* transcript, suggesting that #1 and the probe used in the Allen Brain Atlas were not specific to *6330403K07Rik/UGS148*. On the other hand, probe #3, which was designed against the 3’ untranslated region of *6330403K07Rik/UGS148,* detected a single band at the expected size, which was not detected in the RNA samples derived from the mutant mouse lacking this gene (Fig. 1F). Unexpectedly, these extra bands were not detected with probe #2 that targeted the region weakly homologous to the repeat (TCTCCA)n (Fig. 1F). These results suggested that probe #1 cross-reacted with unknown transcripts via 5’ regions of these genes containing the ORF, even though the sequences were unique in the genome, as shown by conventional bioinformatic searches such as BLAST (Altschul et al., 1990), BLAT (Kent, 2002), or RepeatMasker (Smit, 1996). We also confirmed the specificity of the probes by *in situ* hybridization. Probe #3 mainly detected the signals enriched in the tanycytes, as described in a previous study (Ma et al., 2015), which were not observed in the brains of the mutant mice (Fig. 1D). These results suggested that RNA probes predicted to be unique by conventional sequence analyses possibly detect other transcripts when used in hybridization-based assays. Hereafter, we call *6330403K07Rik Tanmp* (<u>ta</u>nycyte-<u>s</u>pecific ER <u>m</u>embrane <u>p</u>rotein) due to the prominent localization in the ER membrane of tanycytes, as described below. To detect the protein product of *Tanmp*, we raised a monoclonal antibody that specifically detected Tanmp at the predicted size of 13 kDa (Fig. 1G). Both Tanmp mRNA and protein were specifically expressed in the brain tissue but not in other tissues in our investigation (Fig. 1H, I).

### Tanmp is expressed in tanycytes and neurons in the hypothalamus

While a previous study reported highly enriched expression of Tanmp in tanycytes (Ma et al., 2015), the molecular properties and physiological functions of this protein remain largely unknown. To obtain further insights into the functions of this protein without known functional domains, we first examined the expression pattern of Tanmp in the hypothalamus in detail. As previously reported, Tanmp was highly expressed in tanycytes located at the ventral region of the 3^rd^ ventricle in the hypothalamus (Tn in Fig. 2A). Distinct signals were also observed in neurons with a large cell body in the lateral hypothalamic nuclei (LH in Fig. 2A). There are at least two types of tanycytes, α- and β-tanycytes, which are located in dorsal and ventral regions of the 3^rd^ ventricle in the hypothalamus (Langlet, 2014; Rodriguez et al., 2019). Each tanycyte is further categorized into dorsally located type 1 and ventrally located type 2, and four types of tanycytes, α1-, α2-, β1-, and β2-tanycytes, have been described and are aligned from dorsal to ventral in order (Langlet, 2014; Rodriguez et al., 2019). Tanmp was highly expressed in all of these tanycytes, except for β2-tanycytes located in the ventral medial eminence that weakly expressed Tanmp (Fig. 2B). Tanmp-expressing tanycytes were distinct from subventricular astrocytes identified by the expression of Gfap, which are dorsally located along the 3^rd^ ventricle (SA, Fig. 2B, D). Superresolution confocal microscopy of Tanmp revealed the fine morphology of tanycytes, consisting of highly branched mesh-like apical branches (AB), elongated oval cell bodies (CB), and basal processes (BP) extending perpendicular to the ventricular surface (Fig. 2C). In the lateral hypothalamic region that contains Tanmp-positive neurons, there are two types of neurons with large cell bodies, hypocretin/orexin-expressing neurons and melanin-concentrating hormone-expressing neurons, which can be identified by the expression of Hcrt and Pmch, respectively (reviewed in Adamantidis and de Lecea, 2008). Simultaneous detection of the mRNA expression of these marker genes and Tanmp revealed that Tanmp was expressed in all of the Hcrt-positive hypocretin/orexin-expressing neurons but not in the Pmch neurons in the lateral hypothalamus (Fig. 2E).

**Figure 2.**
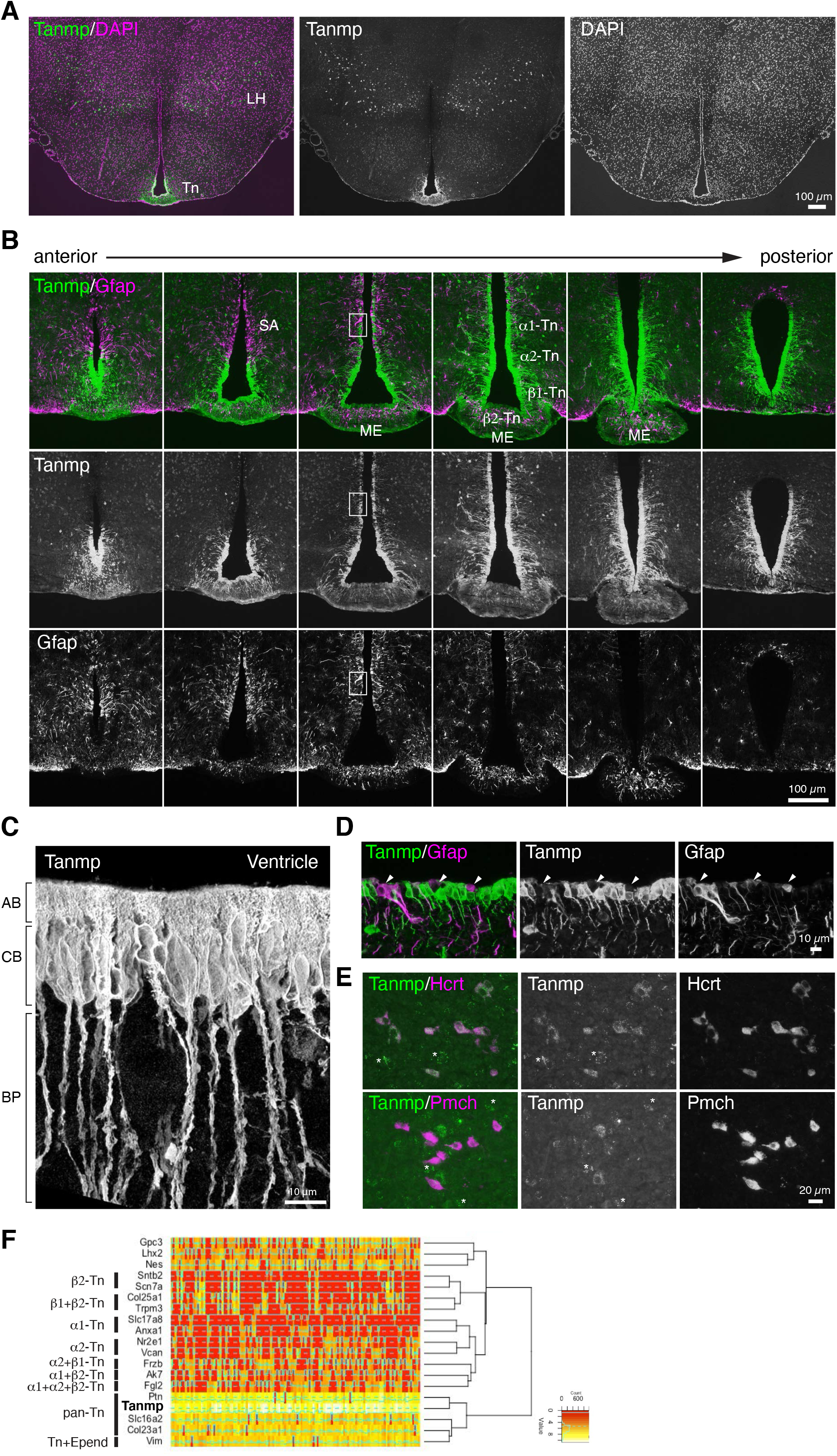
Tanmp is expressed in tanycytes and orexin neurons in the hypothalamus. (A) Coronal section of brain at the level of hypothalamus stained with an antibody against Tanmp (green). Nuclei were counterstained with DAPI and pseudocolored in magenta. Tanmp is expressed in tanycytes (Tn) and neurons with large cell bodies in the lateral hypothalamus (LH). (B) Simultaneous fluorescent detection of Tanmp (green) and GFAP (magenta) in serial coronal sections at the level of the hypothalamus flanking the medial eminence (ME). White boxes indicate the regions shown at higher magnification in (D). Tanmp is strongly expressed in α1-, α2-, and β1-tanycytes (Tn), and weak expression was observed in β2-tanycytes located in the ME. (C) Fine morphology of tanycytes visualized with Tanmp antibody and observed with the Airyscan superresolution confocal microscope. Note the highly branched mesh-like structures in the apical branches (AB). CB: cell bodies, BP: basal processes. (D) Higher magnification of the ventricular region shown in (B). Ventricular astrocytes strongly expressing GFAP (magenta, arrowheads) did not express Tanmp (green). (E) Simultaneous detection of Tanmp (green) and Hcrt or Pmch (magenta) in the lateral hypothalamus. Tanmp is expressed in Hcrt-expressing neurons but not in Pmch-expressing neurons. Asterisks show autofluorescent signals derived from lipofuscin. (F) Hierarchical clustering analyses and heatmap of single-cell RNA sequencing data of the hypothalamus. Tanmp was clustered into a branch containing two pan-tanycyte markers, Slc16a2 and Col23a1. Red-yellow represents relative expression levels, and green lines represent count values.

To further confirm the expression of Tanmp in tanycytes at the single-cell level, we reanalyzed RNA sequencing data prepared from hypothalamic cells (Chen et al., 2017). Hierarchical clustering analyses revealed that Tanmp clustered into genes that are known to be pan-tanycyte marker genes (Fig. 2F), suggesting that Tanmp is ubiquitously expressed in tanycytes at least at the mRNA level.

### Tanmp is an ER membrane protein with an intrinsically disordered domain that protrudes into the cytoplasm

To study the subcellular localization of Tanmp, we exogenously expressed Tanmp in HeLa cells and compared the signals with subcellular organelle markers. Analyses of the primary amino acid sequences using TMHMM (Krogh et al., 2001) revealed that Tanmp is a single transmembrane protein consisting of an N’-located disordered region that contains alternate stretches of basic and acidic amino acids, with a short C’ region consisting of the tripeptide QLA (Fig. 3A, B). ClustalW analyses (Thompson et al., 1994) revealed that the transmembrane domain is highly conserved among mice, rats, and humans, while the N’-located disordered region was rather variable (Fig. 3A). When expressed in HeLa cells, Tanmp was localized in the cytoplasm, forming a mesh-like structure, which resembled the distribution of ER membranes (Fig. 3C). We thus cointroduced GFP-calreticulin, a marker for the ER membrane (Michalak et al., 1999), with Tanmp in HeLa cells. The two signals completely overlapped (Fig. 3C, D), suggesting that Tanmp was localized to the ER membrane. The overall morphology of the ER membrane network was not affected by the overexpression of Tanmp in HeLa cells, and a similar mesh-like pattern was observed in both the Tanmp-expressing and nonexpressing cells (Fig. 3E). We also performed wndchrm analysis (Shamir et al., 2008), a machine-learning-based image recognition analysis, and found that the ER membrane pattern of the parental HeLa cells was indistinguishable from that of the HeLa cells that ectopically expressed Tanmp (Fig. 3F). The ER localization of Tanmp in tanycytes was also confirmed by simultaneous detection of Tanmp and another authentic ER membrane marker, Hspa5/BiP (Munro and Pelham, 1987), in the adult brain, although the signals were obscure due to the harsh antigen retrieval procedure required for the detection of Hspa5/BiP signals in mouse tissue samples (Fig. 3G, H).

**Figure 3.**
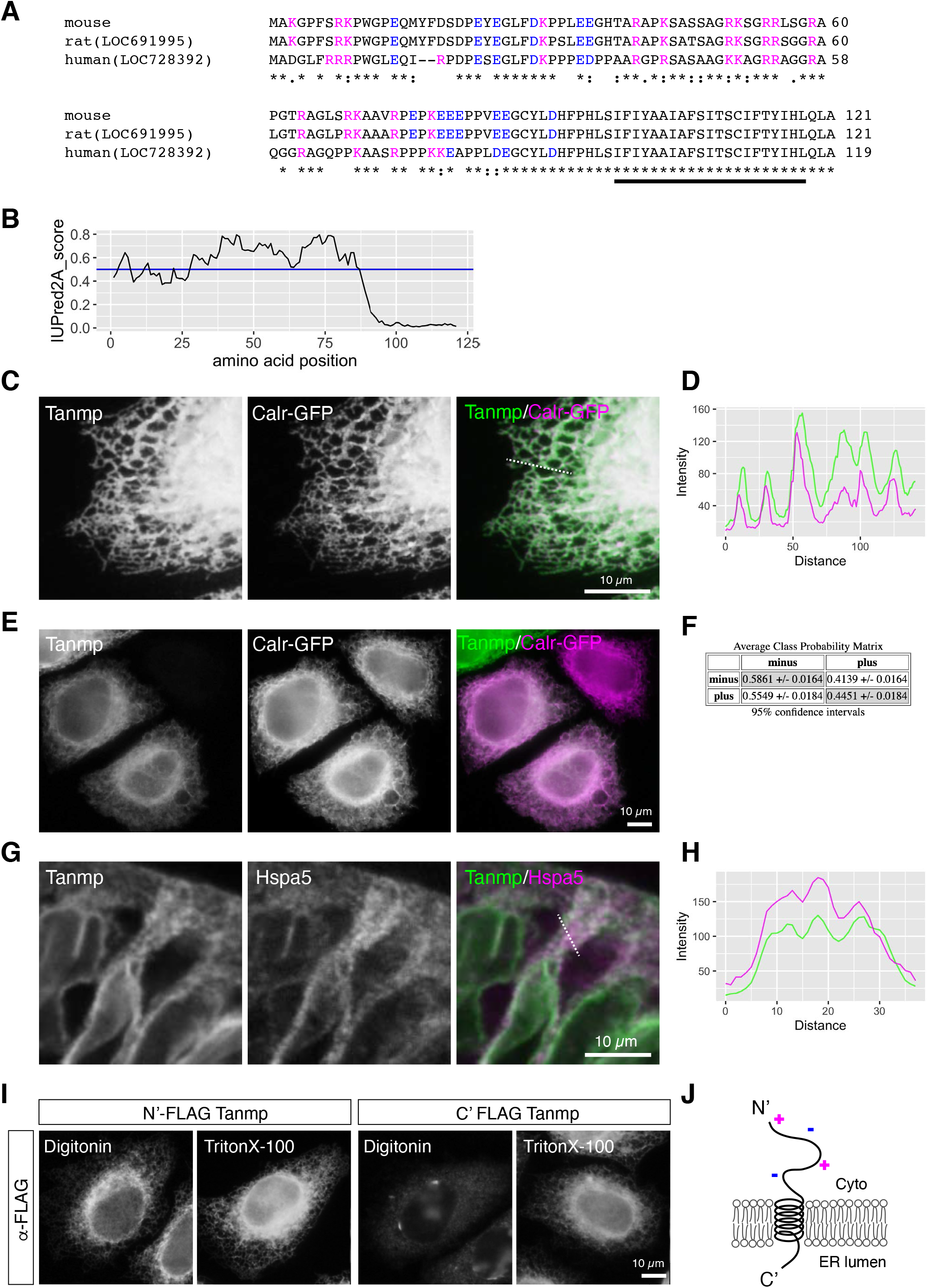
Tanmp is an ER membrane protein with disordered regions. (A) Multiple sequence alignment of mouse, rat, and human Tanmp made by ClustalW. Asterisks indicate conserved amino acids, and colon marks indicate similar amino acids. Acidic and basic amino acids are indicated in blue and magenta, respectively. The bold bar indicates the transmembrane domain predicted by TMHMM. (B) Analysis of the disorder propensity of Tanmp using IUPred2A. The blue line indicates an arbitrary threshold for the intrinsically disordered regions. Note that most of the N’-located nontransmembrane regions are intrinsically disordered. (C) Simultaneous detection of exogenously introduced Tanmp (green) and calreticulin-GFP (magenta) in HeLa cells. Dotted line indicates the region analyzed in (D). (D) Intensity profile plot of Tanmp (green) and calreticulin-GFP (magenta) shown in (C). Note that the peaks of these two signals completely coincided. (E) Morphology of ER membranes shown by calretinin-EGFP (magenta) in Tanmp-expressing (green) and nonexpressing cells. Note that no overt morphological changes were induced upon the expression of Tanmp. (F) Wndchrm analyses of ER morphology of the cells that do (plus) and do not (minus) express Tanmp. Average class probability matrix index indicates probability of the images classified into each category. The images were equally classified into either of the category, suggesting that they were not distinguishable by machine-learning-based image recognition. (G) Simultaneous detection of endogenous Tanmp (green) and the ER marker Hspa5 (magenta) using an Airyscan superresolution confocal microscope. Dotted lines indicate the region analyzed in (H). (H) Intensity profile plot of Tanmp (green) and Hspa5 (magenta) shown in (G). (I) Detection of N’- and C’-tagged FLAG epitopes in HeLa cells permeabilized with digitonin or Triton X-100. Note that C’-tagged FLAG was detected only when the cells were permeabilized with Triton X-100. (J) Schematic drawing of the topology of Tanmp in relation to the ER membrane. The N-terminal intrinsically disordered region is located in the cytoplasmic compartment.

To further investigate the topological organization of Tanmp in the ER membrane, we introduced N’- or C’-tagged Tanmp into HeLa cells and detected the signals after permeabilization using digitonin, which can permeabilize plasma membranes while leaving ER membranes intact (Plutner et al., 1992; Wilson et al., 1995). An N’-tagged FLAG epitope was detected in the digitonin-treated cells, whereas a C’-tagged FLAG epitope could not be detected in the cells (Fig. 3I). However, both epitopes were detected in the cells treated with Triton X-100 (Fig. 3I), which can permeabilize both plasma and ER membranes. These results suggested that the N’ disordered region of Tanmp faces the cytoplasmic regions, whereas the short tripeptide faces the ER lumen (Fig. 3J).

### Tanmp associates with the mitochondrial ATPase subunit, at least in the cell lysate

To gain further insight into the molecular function of Tanmp, we tried to identify molecules that associate with Tanmp in the cells. We established HeLa cells that stably express N’-FLAG-tagged Tanmp and performed immunoprecipitation using cell lysates prepared from the stable cell line. Discrete bands were specifically immunoprecipitated with N’-FLAG-tagged Tanmp (Fig. 4A) and were identified as subunits of mitochondrial ATPase by subsequent MALDI-TOF analyses. To confirm that Tanmp forms a complex with mitochondrial ATPase in the lysate, we performed Western blot analyses of the immunoprecipitated complex using specific antibodies against the α-, β-, and γ-subunits of ATPase. All three subunits were specifically immunoprecipitated with N’-FLAG Tanmp (Fig. 4B), suggesting that Tanmp can associate with the ATPase complex, at least in the cell lysate. Interestingly, multiple bands were detected for FLAG-Tanmp (arrowheads in the FLAG panel in Fig. 4B), suggesting that FLAG-Tanmp forms SDS-resistant multimers during immunoprecipitation.

**Figure 4.**
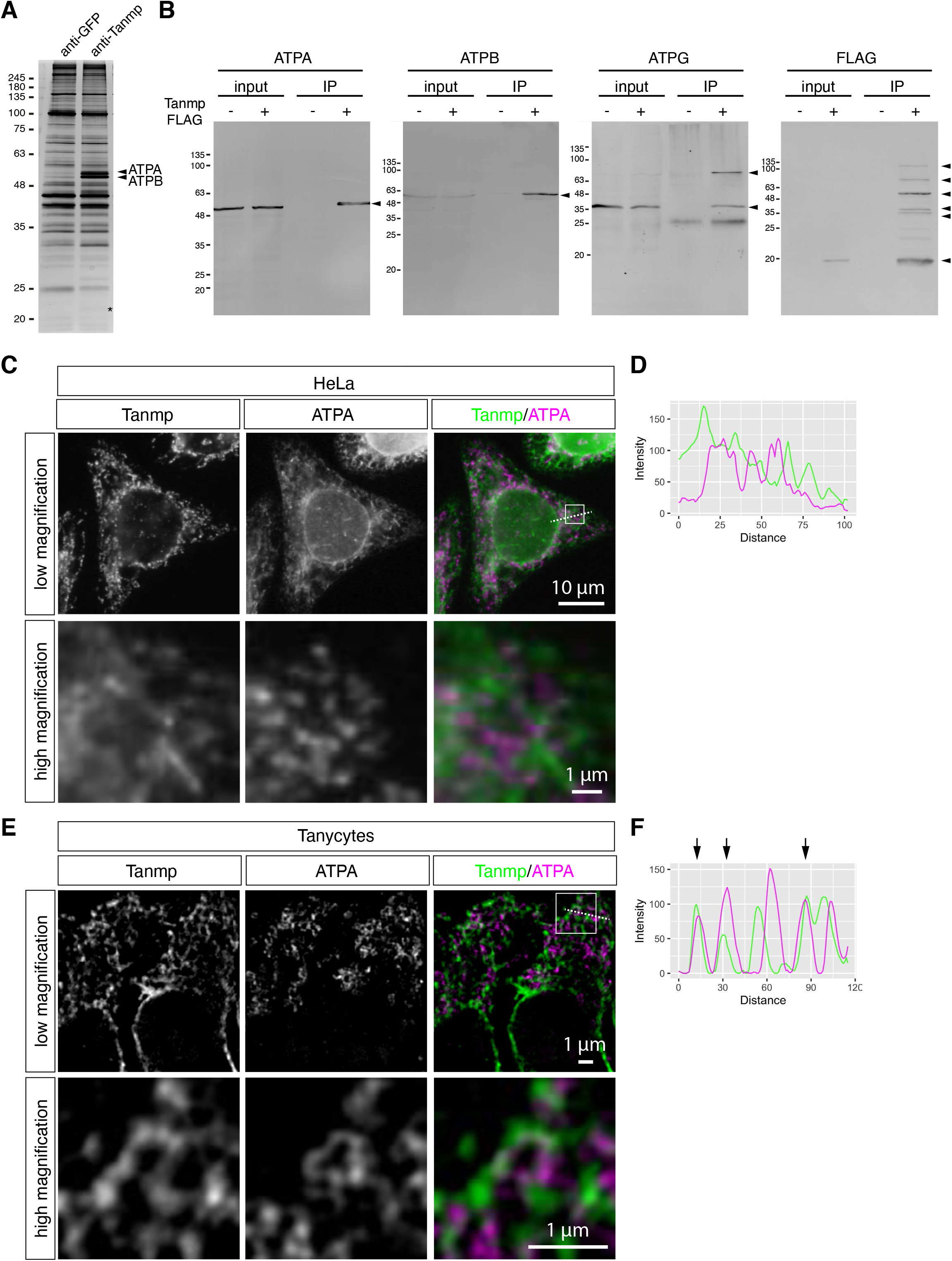
Tanmp associates with the mitochondrial ATPase complex. (A) Silver staining of immunoprecipitated samples run on SDS-PAGE. Tanmp was specifically coimmunoprecipitated with multiple bands at approximately 50 kDa, which were determined to be ATPase complexes by MALDI-TOF analyses. (B) Western blot analyses of proteins coimmunoprecipitated with FLAG-tagged Tanmp detected with specific antibodies against the α-(ATPA), β-(ATPB), and γ-(ATPG) subunits of ATPase. Arrowheads indicate coimmunoprecipitated proteins detected by Western blot. Note that multiple bands were detected for FLAG-Tanmp. (C) Simultaneous detection of Tanmp and ATPA in HeLa cells that exogenously express Tanmp. The signals were observed under a conventional fluorescence microscope. Note that the two signals did not largely overlap. The white box indicates the regions shown at higher magnification in the lower panels. Dotted line indicates the region analyzed in (D). (D) Intensity profile plot of Tanmp (green) and ATPA (magenta). Note that the peaks of these two signals largely do not coincide. (E) Simultaneous detection of Tanmp and ATPA in tanycytes. The signals were observed with a structural illumination superresolution microscope. The white box indicates the regions shown at higher magnification in the lower panels. The dotted line indicates the region analyzed in (F). (F) Intensity profile plot of Tanmp (green) and ATPA (magenta). Arrows indicate overlapping peaks of Tanmp and ATPA signals.

Because Tanmp was localized in the ER membrane, association with mitochondrial ATPases that are localized in the inner membrane of mitochondria was unexpected. We thus simultaneously detected Tanmp and the α-subunit of the ATPase subunit in HeLa cells that stably expressed Tanmp, and the two signals basically did not overlap, as expected (Fig. 4C, D). We also examined the localization of endogenous Tanmp and the α-subunit of ATPase in tanycytes. Although the Tanmp signals did not coincide with the signals of the α-subunit of ATPase in most cases, we occasionally observed Tanmp signals aligning along the mitochondrial signals (Fig. 4E, F).

### Tanmp is not essential for mice reared under laboratory conditions

Finally, we examined the physiological function of Tanmp by creating KO mice that lack the expression of this molecule (Fig. 5A-C). Although the number of adult (8 weeks) KO mice obtained by crossing the heterozygous mice was slightly lower than the expected number, the difference was not significant (male: P = 0.088, female: P = 0.055, chi-square test) (Fig. 5D). Both female and male adult KO mice exhibited similar body weights compared to their wild-type (WT) littermates (Fig. 5E). No overt morphological abnormality of the tanycyte-containing hypothalamus regions was observed on hematoxylin-eosin-stained paraffin sections (Fig. 5F). We then examined the morphology of tanycytes using the ER membrane marker Hspa5/BiP, and mesh-like structures of the AB and BP were normally detected in the KO mice (Fig. 5G). We also examined the differentiation of tanycyte- and hypocretin/orexin-expressing neurons using molecular markers. All of these tanycyte and hypocretin neuron markers were normally expressed in the KO mouse brain (Fig. 5H), suggesting that Tanmp is not involved in the differentiation of these cell types, at least in the animals reared in the laboratory conditions.

**Figure 5.**
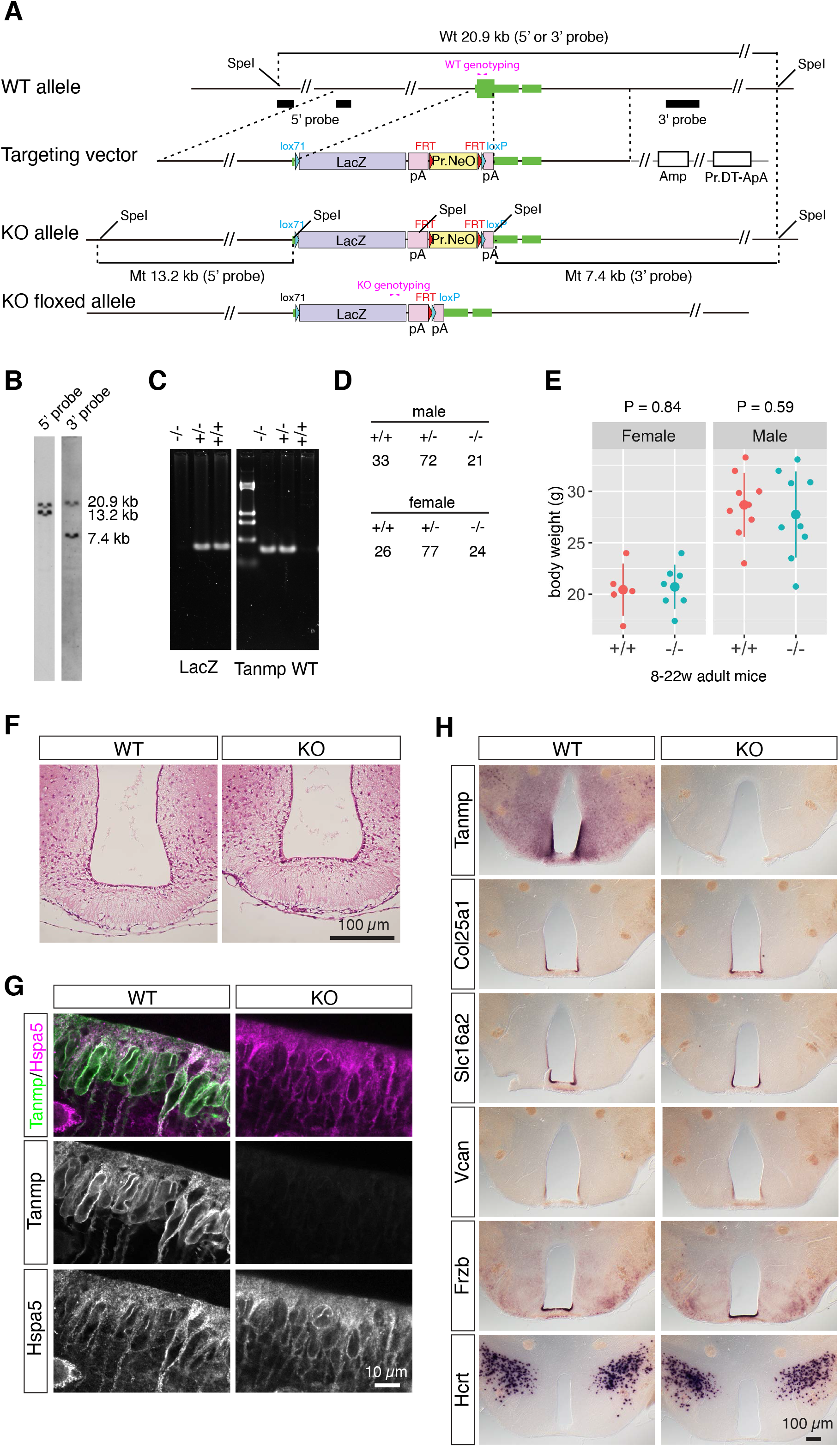
Genetic deletion of Tanmp does not cause overt phenotypic changes in tanycytes. (A) Targeting strategy for disruption of Tanmp. The open reading frame of Tanmp was replaced with the LacZ gene followed by polyadenylation signals. The founder heterozygous mice were crossed with mice expressing Flp recombinase to flip out the Neo cassette, resulting in the generation of floxed Tanmp KO mice, which were used for subsequent analysis. SpeI restriction sites used for the Southern blot analyses are indicated in the short lines. (B) Southern blot analysis of SpeI-digested DNA from ES cells that undergo homologous recombination. The positions of the 5’- and 3’-probes and primers used for genotyping are indicated by black bars and magenta arrows, respectively, in (A). For the 5’ probe, two probes were mixed for sensitive detection. (C) Agarose gel electrophoresis of PCR fragments amplified using primers for genotyping. The molecular markers are pUC19/HinfI-digested DNA. (D) The number of littermates obtained by crossing heterozygous Tanmp mutant mice. Wild-type (+/+), heterozygous (+/−), and homozygous (−/−) mice were obtained at nearly the expected rate. (E) Body weight of 8- to 22-week-old male and female littermates. Each dot represents individual animals, and the vertical line represents standard deviations. P values were calculated by Welch’s t-test. (F) Coronal paraffin sections of adult brain at the level of tanycytes stained with hematoxylin-eosin. Note that no overt phenotype was observed in the KO mice. (G) Simultaneous detection of Tanmp and Hspa5 in tanycytes of the WT and KO mice. Brain slices were stained with the antibodies, and the signals were detected using an Airyscan superresolution confocal microscope. (H) Expression patterns of tanycyte markers (Col25a1, Slc16a2, Vcan, Frzb) and hypocretin/orexin-expressing neurons (Hcrt) in the WT and KO mice.

## Discussion

We have thus identified Tanmp as an ER membrane protein specifically expressed in brain tissue. The strongest expression of Tanmp was observed in tanycytes that align the ventral region of the 3^rd^ ventricle in the hypothalamus, and antibody staining of Tanmp visualized the fine morphology of tanycytes with well-developed ER networks located apically. Given that tanycytes regulate molecular transport between the cerebrospinal fluid and hormone-secreting neurons in the ventral hypothalamus (reviewed in Langlet, 2014; Rodriguez et al., 2019), Tanmp may regulate molecular trafficking across the tanycyte through the characteristic ER membranes. However, the KO mice lacking Tanmp expression did not exhibit obvious changes in the ER membrane morphology of tanycytes, and tanycyte-specific molecular markers were normally expressed in the mutant mice. In addition, the body weights of the Tanmp KO mice were indistinguishable from those of the WT mice, suggesting that the proposed functions of tanycytes, e.g., regulation of the energy balance as a component of the hypothalamic network, are maintained in the Tanmp KO mice. While Tanmp is not essential in mice reared under normal laboratory conditions, this finding does not exclude the possibility that Tanmp becomes functional when animals are placed under certain stressed conditions. Furthermore, certain single-transmembrane ER proteins may compensate for the lack of Tanmp in tanycytes and redundantly function in the tanycytes of the Tanmp KO mice. Notably, that the N’ region of Tanmp is composed of amino acid sequences that are predicted to be intrinsically disordered. Moreover, the functions of certain disordered proteins are maintained even after random shuffling of the primary amino acid sequence (Tsuboyama et al., 2020), suggesting that the composition of the amino acid, rather than the primary sequence, is important for the function of disordered proteins that do not have rigid 3-dimensional structures. A conventional homology search based on the conservation of the primary amino acid sequences may thus fail to find family proteins that redundantly function with Tanmp.

Recently, much attention has been given to the properties of proteins containing intrinsically disordered regions, which mediate multivalent weak molecular interactions (reviewed in Alberti et al., 2019; McSwiggen et al., 2019; Peng and Weber, 2019; Roden and Gladfelter, 2021; Shin and Brangwynne, 2017). Intrinsically disordered regions undergo liquid-liquid phase separation *in vitro* and are suggested to be involved in the formation of various molecular condensates in the cells, including nonmembranous organelles, transcriptional complexes, organizing sites of autophagosomes, and disease-related insoluble aggregates (reviewed in Alberti et al., 2019; Roden and Gladfelter, 2021; Shin and Brangwynne, 2017). The N’ region of Tanmp may thus be involved in the formation of certain molecular condensates at the surface of the ER membrane and promote efficient membrane trafficking or molecular transport. Understanding the molecular rules that specify the function of disordered regions of proteins (Wang et al., 2018) or lncRNAs (Lubelsky and Ulitsky, 2018; Pyfrom et al., 2019) may lead to the discovery of hidden gene families that escape classic bioinformatic approaches based on the primary sequence homologies.

Many studies have revealed functional interactions between mitochondrial and ER membranes, especially through the contact site called the mitochondria-associated ER membrane (MAM). The MAM regulates fission and fusion of mitochondria as well as molecular changes between the two organelles through a set of proteins that localizes to the contact sites (reviewed in Petrungaro and Kornmann, 2019; van der Bliek et al., 2013). We have shown that Tanmp interacts with all three subunits of mitochondrial ATP synthase at least *in vitro*, suggesting that Tanmp also functions as an MAM protein to regulate mitochondrial functions. ATP synthetase is normally found in the inner membranes of mitochondria and is topologically separated from ER membranes by the outer membranes of mitochondria in general. However, early electron microscopic studies have shown that the ER membrane occasionally exhibits continuity with tubules protruding from the mitochondrial outer membranes (Franke and Kartenbeck, 1971; Spacek and Lieberman, 1980), which are commonly found in all tanycyte subtypes (Rodriguez et al., 2019). Indeed, we occasionally observed colocalization of Tanmp and mitochondrial markers in tanycytes, although the frequency was low compared to that of other MAM proteins. To clarify the possible physiological importance of the interaction of Tanmp with ATP synthetase, researchers should further dissect the precise molecular mechanism and cellular conditions that lead to the interaction of topologically separated molecules.

The probes we initially used to detect Tanmp mRNA strongly cross-reacted with certain RNAs that could not be identified by BLAST search or other bioinformatic methods available to date to the best of our knowledge. Hybridization-based detection of target RNAs is a fundamental tool for the study of RNA localization within cells or tissues, and databases of systematic genome-wide analyses of RNA localization (Lecuyer et al., 2007; Sunkin et al., 2013; Yokoyama et al., 2009) provide highly useful platforms to investigate the expression pattern of certain RNAs of interest *in silico*. However, the specificity of RNA probes during hybridization-based detection might be regulated by various factors in addition to conventional base pairing, including secondary structures, the formation of non-Watson-Crick base pairs, and possibly by interactions with particular proteins. Indeed, we previously found that probes targeting a specific mRNA strongly cross-reacted through stretches of 22 nucleotides (nt) with 2 nt mismatches even under stringent hybridization conditions (Ishida et al., 2015). Therefore, the specificity of RNA probes used in localization studies, such as *in situ* hybridization, must be experimentally validated using multiple approaches. Northern hybridization is a classic method for quantification of specific RNA transcripts (Alwine et al., 1977) but still serves as the only experimental approach that can experimentally estimate the size of the target transcripts, which is particularly useful to confirm the specificity of the probes. The importance of multiple approaches to confirm the specificity of hybridization probes must not be underestimated even in the era of genome-wide high-throughput studies.

## Material and Methods

### Animals

All experiments were approved by the safety division of Hokkaido University (#2015-079). C57BL6/N mice were used for all experiments except if otherwise mentioned. For anesthetization, medetomidine-midazolam-butorphanol (Kawai et al., 2011) was intraperitoneally injected at a volume of 10 µl/g of body weight.

### cDNA cloning, plasmid construction and cell lines

The coding sequence of Tanmp was amplified by PCR using a BAC clone (RP23-479A13) as a template and cloned into pT2K XIGΔin (Urasaki et al., 2006). The sequence of calreticulin was obtained by RT-PCR using a cDNA template prepared from SK-N-SH cells. For generation of pT2KXIGΔin avGFP_2A_FLAG-Tanmp and pT2KXIGΔin avGFP_2A_Tanmp-FLAG, the synthesized ORF region was purchased from gBlocks Gene Fragments (IDT) and cloned into pT2K XIGΔin, which was used to establish stable Neuro2A cell lines used in the coimmunoprecipitation experiments. All constructs were assembled using Gibson Assembly (#E2611L, NEB), and the sequences of inserted cDNA were confirmed by Sanger sequencing. For plasmid transfection, expression vectors were mixed with FuGENE HD transfection reagent (#E2311, Promega) at a ratio of 1:3 and incubated with target cells, following the manufacturer’s instructions.

### Immunohistochemistry

For immunohistochemistry of the cultured cells, cells were cultured on coverslips coated with 0.5 mg/ml poly-L-lysine (Sigma). Cells were fixed with 4% paraformaldehyde (PFA) in HEPES-buffered saline [HBSS: 10 mM HEPES (pH 7.4), 137 mM NaCl, 5.4 mM KCl, 0.34 mM Na_2_HPO_4_, 5.6 mM glucose, 1 mM CaCl_2_, 1 mM MgCl_2_] for 20 minutes at room temperature and permeabilized with 0.5% Triton X-100 in PBS for 5 minutes at room temperature. To selectively permeabilize the plasma membrane, we treated the cells in digitonin buffer [10 mM PIPES (pH 6.8), 1 mM EDTA, 0.1 M KCl, 25 mM MgCl2, 0.3 M sucrose, 0.01% digitonin] for 5 minutes on ice. To prepare sections of mouse brains, we perfused anesthetized mice with 4% PFA in HBSS. Dissected brains were washed once with HBSS, embedded in OCT compound (Sakura Finetech), and frozen on dry ice-ethanol. Frozen samples were sectioned at a thickness of 8 µm using a cryostat (Zeiss, HM520), collected on PLL-coated slide glasses (Matsunami), and fixed in 4% PFA in HBSS for 1 hour at room temperature. For permeabilization, slices were washed three times in HBSS and incubated in 100% methanol for 5 minutes at −20°C. Nonspecific binding was blocked in blocking buffer containing 4% skim milk (Difco) in TBST (50 mM Tris-HCl, 150 mM NaCl, 0.01% Tween 20) for 5 minutes. The samples were incubated in the primary and secondary antibodies diluted in blocking buffer for 1 hour each, followed by 3 washes in TBST. After the final washing, samples were mounted with 97% 2,2-thiodiethanol mounting media and PVA-based mounting media when using Cy2 and Alexa 488, respectively, as described previously (Mito et al., 2016). The list of antibodies used in this study is shown in Supplemental Table T1. Fluorescence and bright field images were obtained using an epifluorescence microscope (BX51; Olympus) equipped with a CCD camera (DP70). For wndchrm analyses, 50 equivalently sized TIFF images (256 × 256 pixels) were obtained using a 40X objective lens and analyzed using wndchrm (Shamir et al., 2008) following the manuals provided in GitHub (https://github.com/wnd-charm/wnd-charm).

### Reanalyzes of single cell RNA-Seq data

Single-cell RNA-Seq data of hypothalamus samples (GSE87544) were downloaded, and 3,319 cells with <2,000 detected genes were selected from 1,443,737 cells using Seurat (Stuart et al., 2019) in R. Hierarchical clustering was performed using hclust in R, and a heatmap was generated using a heatmap in Rusing a set of genes described as tanycyte markers in the original paper (Chen et al., 2017).

### Northern blotting

Total RNA was extracted from cultured cells using TRIzol reagent (#15596026, Thermo Fisher Scientific) according to the manufacturer’s instructions. For purification of RNAs from tissues, three volumes of TRIzol LS (#10296028, Thermo Fisher Scientific) were added to 1 volume of tissue samples, and tissues were homogenized using Omni THQ (Omni International). Ten micrograms of total RNA was mixed with the same volume of Gel Loading Buffer II (AM8546G, Thermo Fisher Scientific) and heated at 65°C for 10 minutes before loading. The RNA samples were separated on a 1% agarose gel (1% agarose gel, 1X MOPS, 10% formaldehyde), washed in distilled water, and treated with 0.05 N NaCl for 20 minutes to enhance transfer efficiency. After equilibration in 20X SSC, RNAs were transferred to a nylon membrane (#11209299001, Sigma-Aldrich) using a standard capillary method and UV-crosslinked using a FUNA UV crosslinker (Funakoshi). Membranes were hybridized with appropriate probes diluted in DIG Easy Hyb (#11603558001, Sigma-Aldrich) at 68°C for 16 hours, washed in 2X SSC at 68°C for 30 minutes, and washed twice in 0.2X SSC at 68°C. Hybridized probes were detected with AP-conjugated anti-DIG antibody (#11093274910, Sigma-Aldrich) and CDP-star (#11685627001, Sigma-Aldrich) following the manufacturer’s instructions. Chemiluminescence signals were detected using a ChemiDoc Touch Imaging System (Bio-Rad).

### Identification of Tanmp-interacting proteins using MALDI-TOF/MS

For extraction of total cell lysates, stable transfectants expressing FLAG-tagged Tanmp were lysed in HBST buffer [0.5% Triton X-100, 150 mM NaCl, 10 mM HEPES buffered with KOH (pH 7.4)] for 30 minutes at 4°C and centrifuged at 17,400 *g* for 5 minutes to remove cell debris. Anti-FLAG M2 antibody affinity gels (#A2220, Sigma-Aldrich) were washed with HBST, added to the supernatant, and incubated for 1 hour at 4°C. After extensive washing with HBST, the proteins bound to the beads were eluted by the addition of FLAG peptide (#F3290, Sigma-Aldrich). Proteins in the eluate were separated by SDS-PAGE to detect Tanmp-interacting proteins by Western blotting or by silver staining. For analysis of the protein sample for MALDI-TOF/MS, a Silver Stain MS kit (#299-58901, Wako Pure Chemicals) was used, and staining and destaining were performed according to the manufacturers’ instructions. After silver staining of the gel, visible bands were cut out, destained and chopped into small pieces. The gel pieces were dehydrated by soaking in acetonitrile and dried using a centrifugal evaporator (CVE-2100, EYELA). The gel pieces were further reacted with reducing solution [50 mM TCEP (Thermo Fisher Scientific), 25 mM NH_4_HCO_3_] at 60°C for ten minutes, followed by washing in washing solution (25 mM NH_4_HCO_3_). The gel pieces were further soaked for 30 minutes in alkylation solution (55 mM 2-iodoacetamide, 25 mM NH_4_HCO_3_) to alkylate the proteins. After alkylation, the gel pieces were heavily dehydrated by soaking in dehydrating solution (50% acetonitrile, 25 mM NH_4_HCO_3_) 3 times and once in acetonitrile. For digestion of the proteins in gels, dehydrated gel pieces were immersed in protease solution (50 mM Tris-HCl (pH 8.5), 10 ng/μl trypsin (#90057, Thermo Fisher Scientific), and 10 ng/μl lysyl endopeptidase (#121-05063, Wako Chemicals)] for 45 minutes on ice. Digested protein fragments were recovered using the extraction solution (50% acetonitrile, 5% TFA) in an ultrasonic bath for 30 minutes. The extracted protein solution was concentrated using a centrifugal evaporator. Extracted protein fragments were spotted onto the standard CHCA matrix for MALDI-TOF/MS on the target plate (MTP AnchorChip TM var/384 TF, Bruker Daltonics), and then, peptides were detected using UltraflexII-18 TOF/TOF (Bruker Daltonics). Identification of proteins was performed by peptide mass fingerprinting using a Mascot search (Matrix Science).

### Western blotting

For preparation of samples for Western blotting, mouse tissues were homogenized in CSK buffer [10 mM PIPES (pH 6.8), 100 mM NaCl, 300 mM sucrose, 3 mM MgCl_2_] and mixed with an equal volume of 2x SDS buffer [0.1 M Tris-HCl (pH 6.8), 4% SDS, 10% 2-mercaptoethanol, 20% glycerol, and 0.1 mM PMSF]. For cultured cells, cells were lysed in RIPA buffer [10 mM Tris-HCl (pH 7.4), 1% NP40, 0.1% sodium deoxycholate, 0.1% SDS, 250 mM NaCl, 1 mM EDTA], and the soluble fraction was mixed with a one-sixth volume of 6x Laemmli buffer [30 mM Tris-HCl (pH 6.8), 6% SDS, 12% 2-mercaptoethanol, 50% glycerol]. Boiled samples were subjected to SDS-PAGE, and separated proteins were transferred to PVDF membranes. Membranes were soaked in blocking solution (4% skim milk in PBST) for 30 minutes and then reacted with appropriate antibodies diluted in PBST. Alexa Fluor 680-conjugated or IRDye800-conjugated antibodies were used as secondary antibodies to detect the bands by an Odyssey Infrared Imaging System (LI-COR).

### Preparation of RNA probes

Primers used to prepare RNA probes are provided in Supplemental Table T1. The target sequence was amplified by PCR and cloned into a plasmid using a TOPO-TA cloning kit (#452640, Thermo Fisher Scientific). DNA fragments containing the T7 promoter sequence were further amplified by PCR and purified using a Wizard SV Gel and PCR clean-up system (#A9281, Promega) to prepare templates for *in vitro* transcription. DIG/FITC-labeled probes were prepared using DIG-(#11277073910, Sigma-Aldrich) or FITC-(#11685619910, Sigma-Aldrich) RNA labeling mix according to the manufacturer’s instructions. Synthesized RNA probes were purified using Centri-Sep Spin Columns (Princeton Separations) or Centri Pure Mini (emp BIOTECH) to remove unincorporated labels and stored at −20°C with an equal volume of formamide.

### *In situ* hybridization of tissue sections and brain slices

*In situ* hybridization was performed as previously described (Mito et al., 2016). All reactions were performed at room temperature otherwise mentioned. Briefly, fresh-frozen tissue sections were fixed in 4% paraformaldehyde in HBSS overnight at 4°C and treated with 0.2 N HCl for 20 minutes and subsequently with 3 μg/ml Proteinase K (#3115887001, Sigma-Aldrich) for 7 minutes at 37°C. The proteinase reaction was stopped in 0.2% glycine in PBS for 10 minutes, and sections were postfixed in 4% paraformaldehyde in HBSS for 20 minutes. After acetylation in acetylation solution (1.5% triethanolamine, 0.25% concentrated HCl, 0.25% acetic anhydride), sections were processed for hybridization. Hybridization was performed in the hybridization solution (50% formamide, 1X Denhardt’s solution, 2X SSC, 10 mM EDTA, 100 μg/ml yeast tRNA, 0.01% Tween-20) for <16 hours at 55°C. Probes were washed twice with 2X SSC containing 50% formamide at 55°C and treated with 10 μg/ml RNase A in RNase A buffer [10 mM Tris-HCl (pH 8.0), 500 mM NaCl, 1 mM EDTA, 0.01% Tween-20] for 1 hour at 37°C. After washes with 2X SSC and 0.2xSSC for 30 minutes at 55°C, the hybridized probes were detected with anti-DIG or anti-FITC antibodies.

For preparation of brain slices, anesthetized mice were perfused with 4% paraformaldehyde, and dissected brains were embedded in 3% agarose. Brain slices at a thickness of 150 μm were prepared using a microslicer (DTK-100N, Dosaka EM), dehydrated in 100% methanol, bleached in a 1:5 mixture of 30% H2O2/methanol for 5 hours, and stored in 100% methanol at −20°C before use. *In situ* hybridization using brain slices was performed according to the whole-mount *in situ* hybridization protocol as described previously (Nakagawa and Takeichi, 1995).

### Production of recombinant proteins and anti-Tanmp monoclonal antibodies

For production of the His-tagged recombinant protein His-Tanmp, the corresponding cDNA was subcloned into the pET-28a vector (Millipore). The recombinant protein was produced in *Escherichia coli* BL21 (DE3) and purified with TALON resin (Clontech). For immunization, 10 μg of His-Tanmp was mixed with the adjuvant TiterMax Gold (TiterMax) to produce antigen-adjuvant emulsions and injected intraperitoneally into four BALB/c female mice every two weeks. The lymphocytes from the immunized mice were fused with myeloma P3U1 cells at a ratio of 3:1 by mixing in 50% polyethylene glycol (Roche). The fused cells were dispersed in 80 ml of GIT medium (Wako, Japan) supplemented with 1 ng/ml IL-6 (PeproTech) and 1×HAT (Kohjin-Bio). The cells were seeded in four 96-well plates at 0.2 ml/well and grown for 10 days at 37°C. The first screening was performed by enzyme-linked immunosorbent assay (ELISA) with 50 ng/well His-Tanmp and subsequently screened with Western blots and immunofluorescence.

### Generation of Tanmp KO mice

Tanmp KO mice were generated following protocols previously described (Murata et al., 2004). Briefly, DNA fragments were amplified by PCR using a BAC clone that covered the Tanmp genomic region and subcloned into pDT-ApA/LacZ/NeO to generate the targeting vector. The linearized targeting vector was electroporated into TT2 ES cells (Yagi et al., 1993), and G418-resistant clones were screened by PCR followed by Southern blot analysis for homologous recombination. Chimeric mice were generated with the recombinant ES clone and mated with C57Bl/6 females to generate heterozygous animals. They were then mated with Gt (ROSA)26Sortm1(FLP1)Dym (Jackson Laboratory) to flip out the PGK-Neo cassette, and the resultant heterozygous mice were maintained under the C57Bl6 genetic background. PCR-mediated genotyping was performed using DNAs obtained from adult or embryonic tails using the following conditions: predenaturation at 96°C for 1 minute, followed by 30 cycles of denaturation at 94°C for 30 seconds, annealing at 62°C for 30 seconds, and extension at 72°C for 30 seconds. The RIKEN accession number of Tanmp KO mice is CDB1164K, and detailed information is available at the following address (http://www2.clst.riken.jp/arg/micelist.html). All animal protocols were approved by the safety division of RIKEN. Primers used to generate the Tanmp KO mice and used for genotyping the mice are provided in Supplemental Table T1.

## Acknowledgments

We thank Dr. Akira Ishizuka for technical support for generating the monoclonal antibody against Tanmp, Dr. Akira Kitamura for kindly providing labeling reagents, Mr. Koichi Fujii for assisting single cell data analyses, and Dr. Mitsunori Fukuda for kindly providing the ER marker antibodies and discussing the work. This work was supported by JSPS KAKENHI Grant Numbers 17H03604, 16H06279, and 16H06276 and a Naito Memorial Foundation Grant granted to S.N.; JSPS KAKENHI Grant Numbers 19K16247 and 19H04889, the Sumitomo Foundation, the Astellas Foundation for Research on Metabolic Disorders, and the Takeda Science Foundation granted to S. Y.; and JSPS KAKENHI Grant Numbers 20H04687 and 20K07028 granted to H. M.

## References

Adamantidis, A. and de Lecea, L. (2008). Physiological arousal: a role for hypothalamic systems. Cell Mol Life Sci 65, 1475–88.

Alberti, S., Gladfelter, A. and Mittag, T. (2019). Considerations and Challenges in Studying Liquid-Liquid Phase Separation and Biomolecular Condensates. Cell 176, 419–434.

Altschul, S. F., Gish, W., Miller, W., Myers, E. W. and Lipman, D. J. (1990). Basic local alignment search tool. J Mol Biol 215, 403–10.

Alwine, J. C., Kemp, D. J. and Stark, G. R. (1977). Method for detection of specific RNAs in agarose gels by transfer to diazobenzyloxymethyl-paper and hybridization with DNA probes. Proc Natl Acad Sci U S A 74, 5350–4.

Anderson, D. M., Anderson, K. M., Chang, C. L., Makarewich, C. A., Nelson, B. R., McAnally, J. R., Kasaragod, P., Shelton, J. M., Liou, J., Bassel-Duby, R. et al. (2015). A micropeptide encoded by a putative long noncoding RNA regulates muscle performance. Cell 160, 595–606.

Carninci, P. Kasukawa, T. Katayama, S. Gough, J. Frith, M. C. Maeda, N. Oyama, R. Ravasi, T. Lenhard, B. Wells, C. et al. (2005). The transcriptional landscape of the mammalian genome. Science 309, 1559–63.

Chen, R., Wu, X., Jiang, L. and Zhang, Y. (2017). Single-Cell RNA-Seq Reveals Hypothalamic Cell Diversity. Cell Rep 18, 3227–3241.

Chng, S. C., Ho, L., Tian, J. and Reversade, B. (2013). ELABELA: a hormone essential for heart development signals via the apelin receptor. Dev Cell 27, 672–80.

D’Lima, N. G., Ma, J., Winkler, L., Chu, Q., Loh, K. H., Corpuz, E. O., Budnik, B. A., Lykke-Andersen, J., Saghatelian, A. and Slavoff, S. A. (2017). A human microprotein that interacts with the mRNA decapping complex. Nat Chem Biol 13, 174–180.

Franke, W. W. and Kartenbeck, J. (1971). Outer mitochondrial membrane continuous with endoplasmic reticulum. Protoplasma 73, 35–41.

Frankish, A., Diekhans, M., Jungreis, I., Lagarde, J., Loveland, J. E., Mudge, J. M., Sisu, C., Wright, J. C., Armstrong, J., Barnes, I. et al. (2021). Gencode 2021. Nucleic Acids Res 49, D916–D923.

Hanyu-Nakamura, K., Sonobe-Nojima, H., Tanigawa, A., Lasko, P. and Nakamura, A. (2008). Drosophila Pgc protein inhibits P-TEFb recruitment to chromatin in primordial germ cells. Nature 451, 730–3.

Hashimoto, T., Horikawa, D. D., Saito, Y., Kuwahara, H., Kozuka-Hata, H., Shin, I. T., Minakuchi, Y., Ohishi, K., Motoyama, A., Aizu, T. et al. (2016). Extremotolerant tardigrade genome and improved radiotolerance of human cultured cells by tardigrade-unique protein. Nat Commun 7, 12808.

Ingolia, N. T. (2014). Ribosome profiling: new views of translation, from single codons to genome scale. Nat Rev Genet 15, 205–13.

Ishida, K., Miyauchi, K., Kimura, Y., Mito, M., Okada, S., Suzuki, T. and Nakagawa, S. (2015). Regulation of gene expression via retrotransposon insertions and the noncoding RNA 4.5S RNAH. Genes Cells 20, 887–901.

Kawai, S., Takagi, Y., Kaneko, S. and Kurosawa, T. (2011). Effect of three types of mixed anesthetic agents alternate to ketamine in mice. Exp Anim 60, 481–7.

Kent, W. J. (2002). BLAT--the BLAST-like alignment tool. Genome Res 12, 656–64.

Kondo, T., Hashimoto, Y., Kato, K., Inagaki, S., Hayashi, S. and Kageyama, Y. (2007). Small peptide regulators of actin-based cell morphogenesis encoded by a polycistronic mRNA. Nat Cell Biol 9, 660–5.

Krogh, A., Larsson, B., von Heijne, G. and Sonnhammer, E. L. (2001). Predicting transmembrane protein topology with a hidden Markov model: application to complete genomes. J Mol Biol 305, 567–80.

Langlet, F. (2014). Tanycytes: a gateway to the metabolic hypothalamus. J Neuroendocrinol 26, 753–60.

Lecuyer, E., Yoshida, H., Parthasarathy, N., Alm, C., Babak, T., Cerovina, T., Hughes, T. R., Tomancak, P. and Krause, H. M. (2007). Global analysis of mRNA localization reveals a prominent role in organizing cellular architecture and function. Cell 131, 174–87.

Lubelsky, Y. and Ulitsky, I. (2018). Sequences enriched in Alu repeats drive nuclear localization of long RNAs in human cells. Nature 555, 107–111.

Ma, M. S., Brouwer, N., Wesseling, E., Raj, D., van der Want, J., Boddeke, E., Balasubramaniyan, V. and Copray, S. (2015). Multipotent stem cell factor UGS148 is a marker for tanycytes in the adult hypothalamus. Mol Cell Neurosci 65, 21–30.

Matsumoto, A., Pasut, A., Matsumoto, M., Yamashita, R., Fung, J., Monteleone, E., Saghatelian, A., Nakayama, K. I., Clohessy, J. G. and Pandolfi, P. P. (2017). mTORC1 and muscle regeneration are regulated by the LINC00961-encoded SPAR polypeptide. Nature 541, 228–232.

McSwiggen, D. T., Mir, M., Darzacq, X. and Tjian, R. (2019). Evaluating phase separation in live cells: diagnosis, caveats, and functional consequences. Genes Dev 33, 1619–1634.

Michalak, M., Corbett, E. F., Mesaeli, N., Nakamura, K. and Opas, M. (1999). Calreticulin: one protein, one gene, many functions. Biochem J 344 Pt 2, 281–92.

Mito, M., Kawaguchi, T., Hirose, T. and Nakagawa, S. (2016). Simultaneous multicolor detection of RNA and proteins using super-resolution microscopy. Methods 98, 158–165.

Munro, S. and Pelham, H. R. (1987). A C-terminal signal prevents secretion of luminal ER proteins. Cell 48, 899–907.

Murata, T., Furushima, K., Hirano, M., Kiyonari, H., Nakamura, M., Suda, Y. and Aizawa, S. (2004). ang is a novel gene expressed in early neuroectoderm, but its null mutant exhibits no obvious phenotype. Gene Expr Patterns 5, 171–8.

Nakagawa, S. and Takeichi, M. (1995). Neural crest cell-cell adhesion controlled by sequential and subpopulation-specific expression of novel cadherins. Development 121, 1321–32.

Nelson, B. R., Makarewich, C. A., Anderson, D. M., Winders, B. R., Troupes, C. D., Wu, F., Reese, A. L., McAnally, J. R., Chen, X., Kavalali, E. T. et al. (2016). A peptide encoded by a transcript annotated as long noncoding RNA enhances SERCA activity in muscle. Science 351, 271–5.

Ng, S. Y., Lin, L., Soh, B. S. and Stanton, L. W. (2013). Long noncoding RNAs in development and disease of the central nervous system. Trends Genet 29, 461–8.

Pauli, A., Norris, M. L., Valen, E., Chew, G. L., Gagnon, J. A., Zimmerman, S., Mitchell, A., Ma, J., Dubrulle, J., Reyon, D. et al. (2014). Toddler: an embryonic signal that promotes cell movement via Apelin receptors. Science 343, 1248636.

Peng, A. and Weber, S. C. (2019). Evidence for and against Liquid-Liquid Phase Separation in the Nucleus. Noncoding RNA 5.

Petrungaro, C. and Kornmann, B. (2019). Lipid exchange at ER-mitochondria contact sites: a puzzle falling into place with quite a few pieces missing. Curr Opin Cell Biol 57, 71–76.

Plutner, H., Davidson, H. W., Saraste, J. and Balch, W. E. (1992). Morphological analysis of protein transport from the ER to Golgi membranes in digitonin-permeabilized cells: role of the P58 containing compartment. J Cell Biol 119, 1097–116.

Pyfrom, S. C., Luo, H. and Payton, J. E. (2019). PLAIDOH: a novel method for functional prediction of long non-coding RNAs identifies cancer-specific LncRNA activities. BMC Genomics 20, 137.

Roden, C. and Gladfelter, A. S. (2021). RNA contributions to the form and function of biomolecular condensates. Nat Rev Mol Cell Biol 22, 183–195.

Rodriguez, E., Guerra, M., Peruzzo, B. and Blazquez, J. L. (2019). Tanycytes: A rich morphological history to underpin future molecular and physiological investigations. J Neuroendocrinol 31, e12690.

Shamir, L., Orlov, N., Eckley, D. M., Macura, T., Johnston, J. and Goldberg, I. G. (2008). Wndchrm - an open source utility for biological image analysis. Source Code Biol Med 3, 13.

Shin, Y. and Brangwynne, C. P. (2017). Liquid phase condensation in cell physiology and disease. Science 357.

Smit, A. F. (1996). The origin of interspersed repeats in the human genome. Curr Opin Genet Dev 6, 743–8.

Spacek, J. and Lieberman, A. R. (1980). Relationships between mitochondrial outer membranes and agranular reticulum in nervous tissue: ultrastructural observations and a new interpretation. J Cell Sci 46, 129–47.

Stuart, T., Butler, A., Hoffman, P., Hafemeister, C., Papalexi, E., Mauck, W. M., 3rd, Hao, Y., Stoeckius, M., Smibert, P. and Satija, R. (2019). Comprehensive Integration of Single-Cell Data. Cell 177, 1888–1902 e21.

Sunkin, S. M., Ng, L., Lau, C., Dolbeare, T., Gilbert, T. L., Thompson, C. L., Hawrylycz, M. and Dang, C. (2013). Allen Brain Atlas: an integrated spatio-temporal portal for exploring the central nervous system. Nucleic Acids Res 41, D996–D1008.

Thompson, J. D., Higgins, D. G. and Gibson, T. J. (1994). CLUSTAL W: improving the sensitivity of progressive multiple sequence alignment through sequence weighting, position-specific gap penalties and weight matrix choice. Nucleic Acids Res 22, 4673–80.

Tsuboyama, K., Osaki, T., Matsuura-Suzuki, E., Kozuka-Hata, H., Okada, Y., Oyama, M., Ikeuchi, Y., Iwasaki, S. and Tomari, Y. (2020). A widespread family of heat-resistant obscure (Hero) proteins protect against protein instability and aggregation. PLoS Biol 18, e3000632.

Ulitsky, I. (2016). Evolution to the rescue: using comparative genomics to understand long non-coding RNAs. Nat Rev Genet 17, 601–14.

Urasaki, A., Morvan, G. and Kawakami, K. (2006). Functional dissection of the Tol2 transposable element identified the minimal cis-sequence and a highly repetitive sequence in the subterminal region essential for transposition. Genetics 174, 639–49.

van der Bliek, A. M., Shen, Q. and Kawajiri, S. (2013). Mechanisms of mitochondrial fission and fusion. Cold Spring Harb Perspect Biol 5.

Wang, J., Choi, J. M., Holehouse, A. S., Lee, H. O., Zhang, X., Jahnel, M., Maharana, S., Lemaitre, R., Pozniakovsky, A., Drechsel, D. et al. (2018). A Molecular Grammar Governing the Driving Forces for Phase Separation of Prion-like RNA Binding Proteins. Cell 174, 688–699 e16.

Wilson, R., Allen, A. J., Oliver, J., Brookman, J. L., High, S. and Bulleid, N. J. (1995). The translocation, folding, assembly and redox-dependent degradation of secretory and membrane proteins in semi-permeabilized mammalian cells. Biochem J 307 (Pt 3), 679–87.

Yagi, T., Tokunaga, T., Furuta, Y., Nada, S., Yoshida, M., Tsukada, T., Saga, Y., Takeda, N., Ikawa, Y. and Aizawa, S. (1993). A novel ES cell line, TT2, with high germline-differentiating potency. Anal Biochem 214, 70–6.

Yokoyama, S., Ito, Y., Ueno-Kudoh, H., Shimizu, H., Uchibe, K., Albini, S., Mitsuoka, K., Miyaki, S., Kiso, M., Nagai, A. et al. (2009). A systems approach reveals that the myogenesis genome network is regulated by the transcriptional repressor RP58. Dev Cell 17, 836–48.

